# Variants of *DNMT3A* cause transcript-specific DNA methylation patterns and affect hematopoietic differentiation

**DOI:** 10.1101/318188

**Authors:** Tanja Božić, Joana Frobel, Annamarija Raic, Fabio Ticconi, Chao-Chung Kuo, Stefanie Heilmann-Heimbach, Tamme W. Goecke, Martin Zenke, Edgar Jost, Ivan G. Costa, Wolfgang Wagner

**Author notes:** Corresponding author:* Wolfgang Wagner, M.D., Ph.D., Helmholtz-Institute for Biomedical Engineering, Stem Cell Biology and Cellular Engineering, RWTH Aachen University Medical School, Pauwelsstraße 20, 52074 Aachen, Germany. These authors contributed equally to this work (Tanja Božić, Joana Frobel).

## Abstract

The *de novo* DNA methyltransferase 3A (DNMT3A) plays pivotal roles in hematopoietic differentiation. In this study, we followed the hypothesis that alternative splicing of *DNMT3A* has characteristic epigenetic and functional sequels. Specific *DNMT3A* transcripts were either downregulated or overexpressed in human hematopoietic stem and progenitor cells and this resulted in complementary and transcript-specific DNA methylation and gene expression changes. Functional analysis indicated that particularly transcript 2 (coding for DNMT3A2) activates proliferation and induces loss of a primitive immunophenotype, whereas transcript 4 interferes with colony formation of the erythroid lineage. Notably, in acute myeloid leukemia (AML) expression of transcript 2 correlates with its *in vitro* DNA methylation and gene expression signatures and is associated with overall survival, indicating that *DNMT3A* variants impact also on malignancies. Our results demonstrate that specific *DNMT3A* variants have distinct epigenetic and functional impact. Particularly DNMT3A2 triggers hematopoietic differentiation and the corresponding signatures are reflected in AML.

## Introduction

DNA methylation (DNAm) of CG dinucleotides (CpGs) is a key epigenetic process in cellular differentiation (Broske et al. 2009). Establishment of new DNAm patterns is particularly mediated by the methyltransferases DNMT3A and DNMT3B (Okano et al. 1999). Both *de novo* DNMTs are subject to extensive tissue- or developmental stage-specific alternative splicing (Weisenberger et al. 2002) and different variants can be co-expressed in the same cell (Van Emburgh and Robertson 2011). There is evidence that alternative splicing of DNMTs affects enzymatic activity or binding specificity (Weisenberger et al. 2002; Choi et al. 2011; Duymich et al. 2016), but so far it is largely unclear if the different variants mediate different DNAm patterns and if they really possess specific functions in development.

DNMT3A is of particular relevance for hematopoietic differentiation. Conditional ablation of exon 19 of *Dnmt3a* in mice was shown to increase the hematopoietic stem cell pool and impairs their differentiation (Challen et al. 2011; 2014). Furthermore, it has been shown that *DNMT3A* is the most frequently mutated gene in clonal hematopoiesis of the elderly and this was linked to a higher risk for hematological malignancies – indicating that aberrations in *DNMT3A* play a central role for clonal hematopoiesis (Genovese et al. 2014; Jaiswal et al. 2014; Xie et al. 2014). However, the functional roles of specific *DNMT3A* variants in hematopoietic differentiation and malignancies have not been systematically compared.

Acute myeloid leukemia (AML) is frequently associated with genomic mutations in *DNMT3A* – either at the highly recurrent position R882 (Yamashita et al. 2010), or at other sites within this gene (Ley et al. 2010; Yoshizato et al. 2015). These mutations are associated with poor prognosis and are used for risk stratification in AML (Ley et al. 2010; Ribeiro et al. 2012). In our previous study, we demonstrated that ~40% of AML patients have an aberrant hypermethylation within the *DNMT3A* gene (Jost et al. 2014). This hypermethylation is also associated with poor prognosis in AML (Jost et al. 2014) and myelodysplastic syndromes (Mies et al. 2016) and was therefore termed “*DNMT3A* epimutation”. Notably, mutations and the “epimutation” of *DNMT3A* resulted in downregulation of exons associated with transcript 2 (coding for DNMT3A2) (Jost et al. 2014). Furthermore, *in vitro* expansion of hematopoietic stem and progenitor cells (HSPCs) resulted in downregulation of *DNMT3A* transcript 2 (Weidner et al. 2013). In this study, we followed the hypothesis that different isoforms of DNMT3A have distinct molecular and functional sequels and thereby affect hematopoietic differentiation and malignancy.

## Results and Discussion

### *DNMT3A* splice variants have transcript-specific DNA methylation signatures

So far, five protein coding transcripts of *DNMT3A* have been described: transcripts 1 and 3 (ENST00000264709.7 and ENST00000321117.9, respectively) have different transcription start sites, but code for the same full-length protein isoform, referred to as DNMT3A1; transcript 2 (ENST00000380746.8) is truncated at the N-terminus, and codes for the protein isoform DNMT3A2; transcript 4 (ENST00000406659.3), coding for DNMT3A4, is truncated at the C-terminus and lacks the catalytic active methyltransferase (MTase) domain, as well as the PWWP (Pro-Trp-Trp-Pro) and ADD (ATRX-DNMT3-DNMT3L) domain that can interact with various binding partners and chromatin modifications (Yang et al. 2015). Recently, an additional transcript 5 was identified (ENST00000402667.1) that encodes for a similar isoform as DNMT3A2, but lacks the 2^nd^ exon of transcript 2. While we were able to amplify transcript 5, this transcript does not have a unique exon for transcript-specific knockdown and therefore it was not considered for further analysis.

Initially, individual transcripts were knocked down (KD) in cord blood-derived CD34^+^ HSPCs with short hairpin RNAs (shRNAs) targeting exon 5 of transcripts 1 and 3 (shRNA Tr.1+3), exon 2 of transcript 2 (shRNA Tr.2), and exon 4 of transcript 4 (shRNA Tr.4; Fig. 1A). As a control we used a shRNA containing a scrambled sequence. Significant knockdown was validated by RT-qPCR with primers targeting transcript-specific exons (Fig. 1B; *n* = 3). Additionally, *DNMT3A* transcripts 1+3, 2, and 4 were cloned into vectors for constitutive overexpression (OE) and delivered by lentiviral infection into additional three replicates of CD34^+^ HSPCs. Efficient overexpression was verified by RT-qPCR (Fig. 1C; *n* = 3).

**Figure 1.**
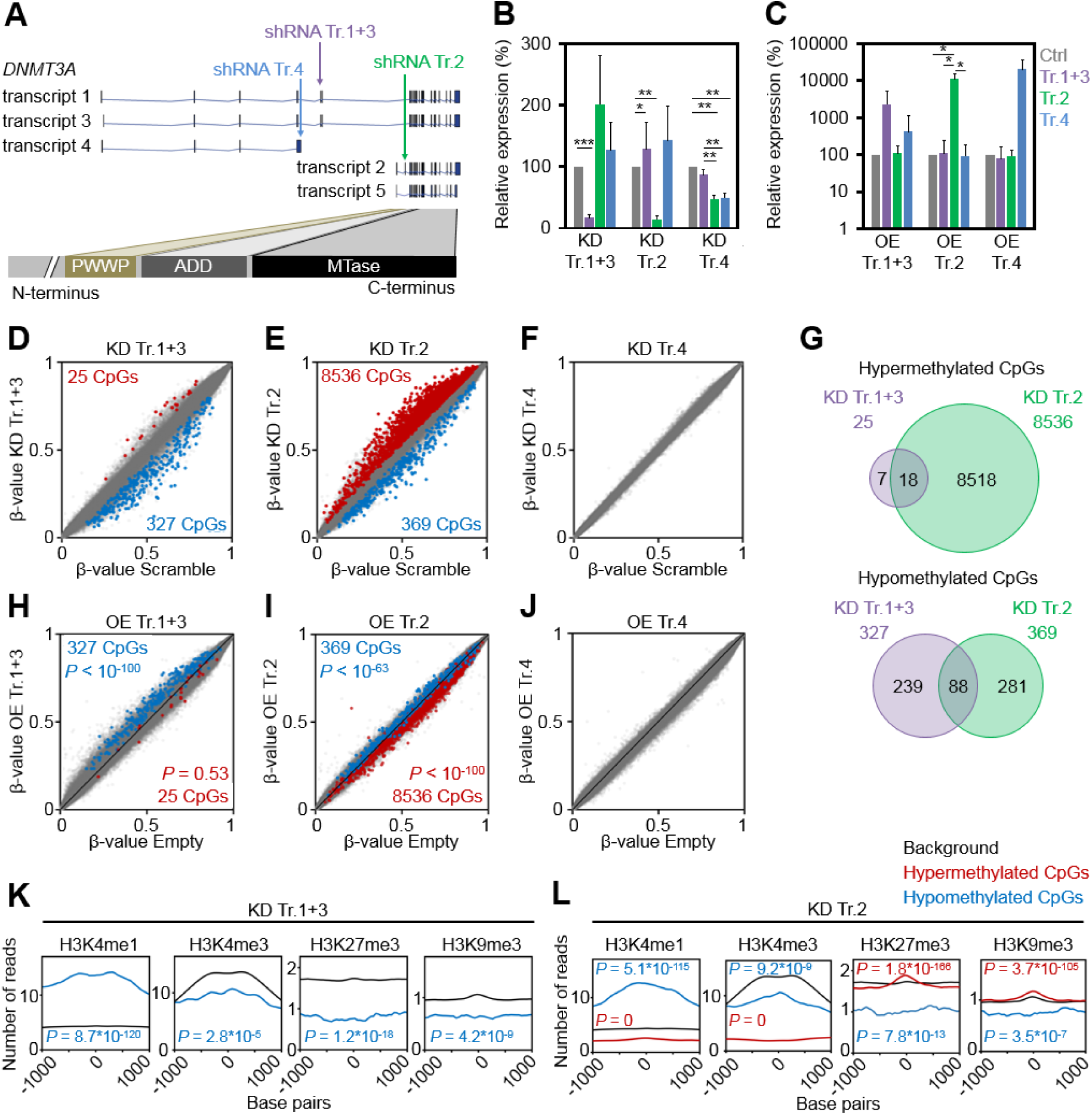
DNMT3A variants cause unique DNA methylation signatures. (*A*) Schematic representation of five protein-coding *DNMT3A* splice variants and their functional protein domains. Target sites of short hairpin RNAs (shRNA) are indicated. (*B, C*) Knockdown (KD; *B*) and overexpression (OE; *C*) of individual transcripts was confirmed with RT-qPCR (mean ± SD; *n* = 3). (*D–F*) Scatter plots of DNAm profiles upon downregulation of transcripts 1+3 (*D*), transcript 2 (*E*), and transcript 4 (*F*) compared to scrambled shRNA controls (mean β- values; *n* = 3). Significantly hyper- and hypomethylated CpGs are depicted in red and blue, respectively (adj. *P* < 0.05). (*G*) Venn diagrams demonstrate overlap of DNAm changes upon modulation of transcripts 1+3 and transcript 2. (*H–J*) Scatter plots of DNAm profiles upon overexpression of transcripts 1+3 (*H*), transcript 2 (*I*), and transcript 4 (*J*), as compared to empty control vectors without any *DNMT3A* transcript (additional independent biological replicates, *n* = 3). CpGs that were significantly hyper- (red) and hypomethylated (blue) upon knockdown of the corresponding transcripts are highlighted. Almost all CpGs are modified in opposite directions upon knockdown and overexpression and the corresponding *P*-values are indicated (Fisher’s *t*-test). (*K,L*) Enrichment of active and repressive histone marks in the vicinity of relevant CpGs with significant DNAm changes upon KD of transcripts 1+3 (*K*) or transcript 2 (*L*). Black lines exemplarily depict 50,000 random CpGs ( background). * *P* < 0.05, ** *P* < 0.01, *** *P* < 0.001 ( Student´s *t*-test).

To investigate if modulation of *DNMT3A* splice variants evokes transcript-specific epigenetic changes we analyzed global DNAm patterns. Knockdown of transcripts 1+3 resulted in 352 CpGs with significant DNAm changes compared to HSPCs infected with the scrambled control (Fig. 1D; Supplemental Table S1A; *n* = 3; adj. *P* < 0.05); knockdown of transcript 2 evoked 8,905 significant DNAm changes (Fig. 1E; Supplemental Table S1B; *n* = 3, adj. *P* < 0.05); whereas knockdown of transcript 4, which does not comprise the MTase domain, did not result in any significant changes (Fig. 1F). Downregulation of transcripts 1+3 resulted preferentially in hypomethylation, whereas downregulation of transcript 2 was rather associated with hypermethylation. The latter is counterintuitive, but in line with preferential hypermethylation in reduced representation bisulfite sequencing (RRBS) data of *Dnmt3a*-null HSCs (Challen et al. 2011). Overall, the overlap of differential DNAm upon KD of transcripts 1+3 and KD of transcript 2 was relatively low and most of the changes seemed to be transcript-specific (Fig. 1G). This is in line with previous reports indicating that DNMT3A1 is associated with heterochromatin, whereas DNMT3A2 is rather associated with euchromatin (Chen et al. 2002), and that the two isoforms have different binding preferences in mouse embryonic stem cells (Manzo et al. 2017). Downregulation of transcript 2 led to a significant hypermethylation of two CpGs within *DNMT3A* (cg20948740 and cg11354105; Δβ-value = 0.0505 and 0.0502, respectively; adj. *P* < 0.05) that are localized close to the epimutation of *DNMT3A,* which is frequently deregulated in AML (Jost et al. 2014). Thus, the *DNMT3A* epimutation may not only result in downregulation of transcript 2 (Jost et al. 2014), but also the other way around. Furthermore, there is recent evidence that DNMT3A itself interacts with splicing factors and impacts global alternative splicing patterns in HSCs (Ramabadran et al. 2017). Such feedback mechanisms can ultimately determine the relative abundance of specific *DNMT3A* transcripts.

Unexpectedly, overexpression of individual transcripts did not entail significant DNAm changes in comparison to controls (*n* = 3, adj. *P* < 0.05). This might be attributed to unaltered endogenous expression of other *DNMT3A* variants. However, when we analyzed those CpGs with significant DNAm changes upon knockdown, we observed that the vast majority of hypomethylated CpGs were now hypermethylated upon overexpression, and *vice versa* (Figs. 1H-J). Statistical analysis (Fisher’s *t*-test) unequivocally demonstrated that downregulation and overexpression of specific *DNMT3A* variants have significant opposing and transcript-specific effects on DNAm patterns (transcripts 1+3: hypomethylated CpGs *P* < 10^−100^; and transcript 2: hypermethylated CpGs *P* < 10^−100^, hypomethylated CpGs *P* < 10^−63^), indicating that the effects of the shRNAs are target specific.

To determine whether targets of *DNMT3A* variants are related to the histone code we utilized chromatin immunoprecipitation sequencing data (ChIP-seq) of short-term cultured human CD34^+^ cells from the International Human Epigenome Consortium (IHEC). Hypomethylation upon either KD of transcripts 1+3 (327 CpGs) or transcript 2 (369 CpGs) was enriched in genomic regions with the histone mark H3K4me1, typically associated with enhancers. In fact, it has recently been shown that hypomethylation in AML is enriched in active enhancer regions marked with H3K4me1 and H3K27ac (Yang et al. 2016; Glass et al. 2017). In contrast, the 8,536 hypermethylated CpGs upon KD of transcript 2 were associated with repressive histone marks H3K27me3 and H3K9me3 (Figs. 1K,L). This is in line with the finding that particularly hypermethylation upon KD of transcript 2 hardly occurred at CGIs and promoter regions (Supplemental Fig. S1A). Similarly, it has been shown that CpGs associated with shore regions, and not CGIs, play a central role for epigenetic classification of AML (Glass et al. 2017). Motive enrichment analysis of differentially methylated regions (DMRs) of DNMT3A1 (flanking 50 bp around each differentially methylated CpG site) revealed a significant enrichment of binding sites for Spi-B transcription factor (SPIB; adj. *P* = 5.5*10^−4^, Supplemental Fig. S1B), whereas DMRs of DNMT3A2 were particularly enriched in binding sites for the hematopoietic transcription factors (TFs) GATA binding protein 4 (GATA4; adj. *P* = 4.0*10^−9^), GATA1 (adj. *P* = 1.5*10^−8^), runt related transcription factor 1 (RUNX1; adj. *P* = 3.6*10^−8^), GATA2 (adj. *P* = 3.6*10^−8^), GATA5 (adj. *P* = 3.6*10^−8^), GATA3 (adj. *P* = 1.8*10 ^−6^), and SRY-box 10 (SOX10; adj. *P* = 2.0*10^−6*^; Supplemental Fig. S1C). Gene Ontology (GO) classification indicated that hypomethylation upon KD of transcripts 1+3 or transcript 2 occurs preferentially in promoter regions of genes associated with lymphocyte activation (particularly of T cells) or immune regulation, respectively (Supplemental Fig. S1D). Recent findings also indicate that *Dnmt3a* is relevant for normal thymocyte maturation (Kramer et al. 2017).

### Transcript-specific DNA methylation changes are reflected in differential gene expression

Next, we analyzed if modulation of *DNMT3A* transcripts resulted in corresponding gene expression changes of HSPCs. Differentially expressed genes were selected by adjusted *P*-values (adj. *P* < 0.05) and with an additional cutoff of >1.5 fold differential expression (*n* = 3). Significant changes were observed in 46 genes upon KD of transcripts 1+3 (Fig. 2A; Supplemental Table S2A), 225 genes upon KD of transcript 2 (Fig. 2B; Supplemental Table S2B), and 190 genes upon KD of transcript 4 (Fig. 2C; Supplemental Table S2C) in comparison to transfection with scrambled shRNA. Overexpression of transcripts 1+3 and transcript 4 did not result in significant differential gene expression (Figs. 2D,F, respectively), which is in line with the moderate DNAm changes. In contrast, overexpression of transcript 2 resulted in significant gene expression changes of 303 genes (adj. *P* < 0.05; Fig. 2E; Supplemental Table S2D). Notably, when we analyzed the 225 significant genes upon knockdown of transcript 2 (155 upregulated and 70 downregulated) we observed that the vast majority was regulated in the opposite direction upon overexpression (Fig. 2G; Fischer’s *t*-test *P* = 6*10^−13^ and *P* = 0.03, respectively), whereas this was less pronounced for other transcripts.

**Figure 2.**
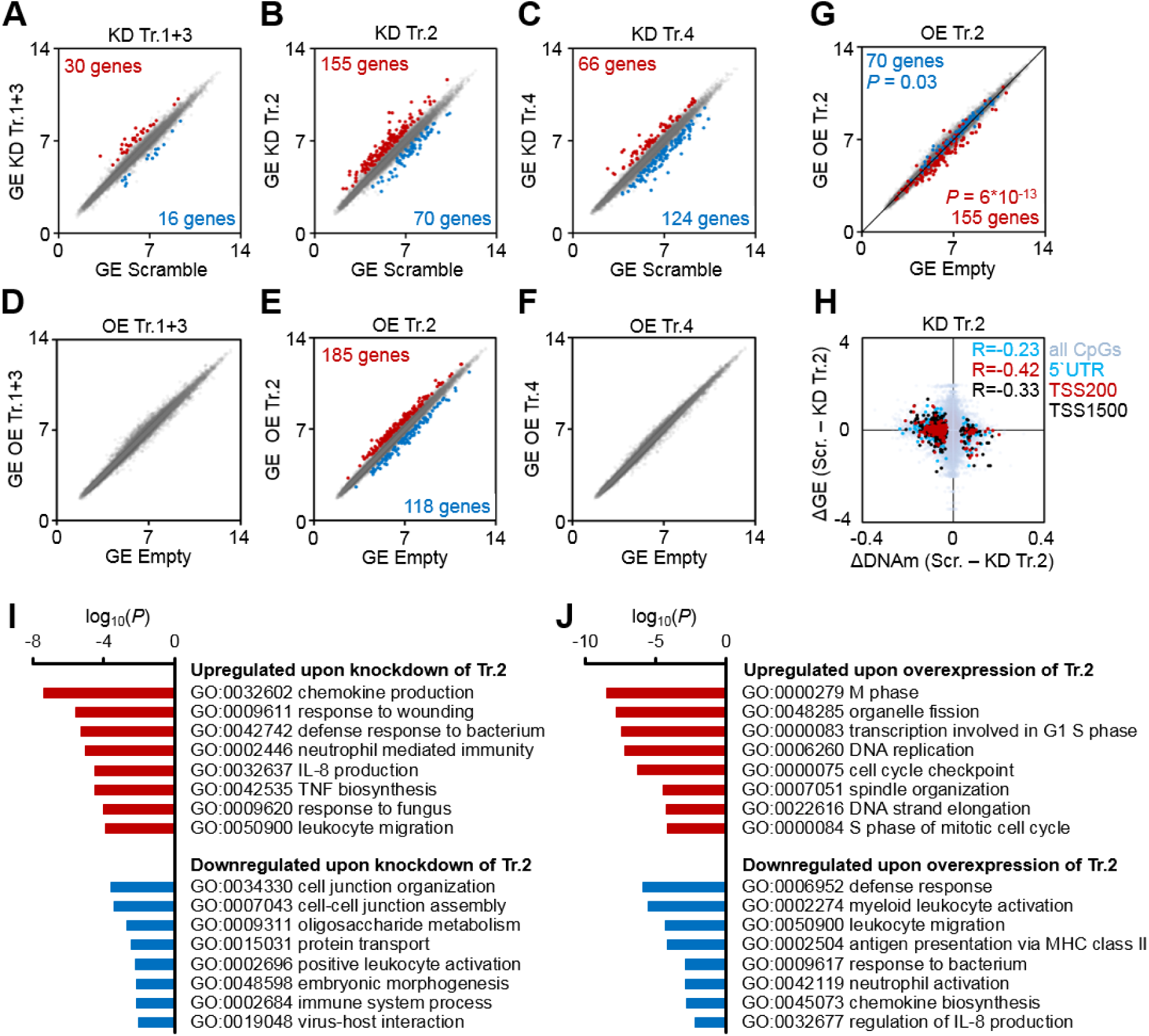
Gene expression changes upon modulation of *DNMT3A* transcripts. (*A–F*) Scatter plots of gene expression (GE; Affymetrix Gene ST 1.0 microarray) upon knockdown (KD; *A–C*) or overexpression (OE; *D–F*) of transcripts 1+3 (*A,D*), transcript 2 (*B,E*), and transcript 4 (*C,F*), as compared to scrambled shRNA controls or empty control vectors (mean log2 values of normalized data; n = 3). Numbers of genes that reached statistical significance (adj. P < 0.05) and at least 1.5-fold up- or down-regulation are indicated in red and blue, respectively. (*G*) To visualize that gene expression changes upon KD and OE of transcript 2 occurred preferentially in opposite directions, the scatter plot depicts significant downregulation (blue) and upregulation (red) upon KD of transcript 2 in data for OE of transcript 2 (Fischer’s t-test). (*H*) DNAm changes of CpGs upon knockdown of transcript 2 were matched to differential expression of corresponding genes. Hypermethylated CpGs in promoter regions were associated with downregulation of gene expression and vice versa (R = −0.23, R = − 0.42 and R = −0.33, respectively; n = 3). (*I,J*) Gene Ontology analysis was performed to classify gene expression changes upon knockdown (*I*) and overexpression (*J*) of transcript 2.

To determine if DNAm changes upon KD of transcript 2 are reflected in corresponding gene expression changes we focused on genes with significant differentially methylated CpGs in the 5´ untranslated region (5´UTR) and up to 200 or 1,500 base pairs upstream of transcription start sites (TSS200 and TSS1500, respectively). As expected, hypermethylation of promoter regions was associated with downregulation of gene expression (Fig. 2H). Furthermore, GO analysis revealed that differentially expressed genes upon either KD or OE of transcript 2 were enriched in complementary categories for up- and downregulated genes (Figs. 2I,J). Genes downregulated by DNMT3A2 were enriched in chemokine production (e.g. interleukin 8), immunity, and leukocyte migration; whereas upregulated genes were involved in proliferation. These results indicate that DNMT3A2-associated DNAm changes are overall reflected by corresponding gene expression changes that are relevant for hematopoietic differentiation.

### DNMT3A transcripts impact on hematopoietic differentiation

Different *DNMT3A* variants might be relevant for proliferation and differentiation of HSPCs. CD34^+^ cells were stained with carboxyfluorescein succinimidyl ester (CFSE) after isolation to estimate their proliferation rate based on residual CFSE after five days post infection (*n* = 3). Proliferation was reduced upon downregulation of transcript 2 (Figs. 3A,B), which is in line with GO enrichment of upregulated genes upon transcript 2 overexpression. In contrast, overexpression of each of the *DNMT3A* transcripts resulted in a moderate increase in proliferation (Fig. 3B).

**Figure 3.**
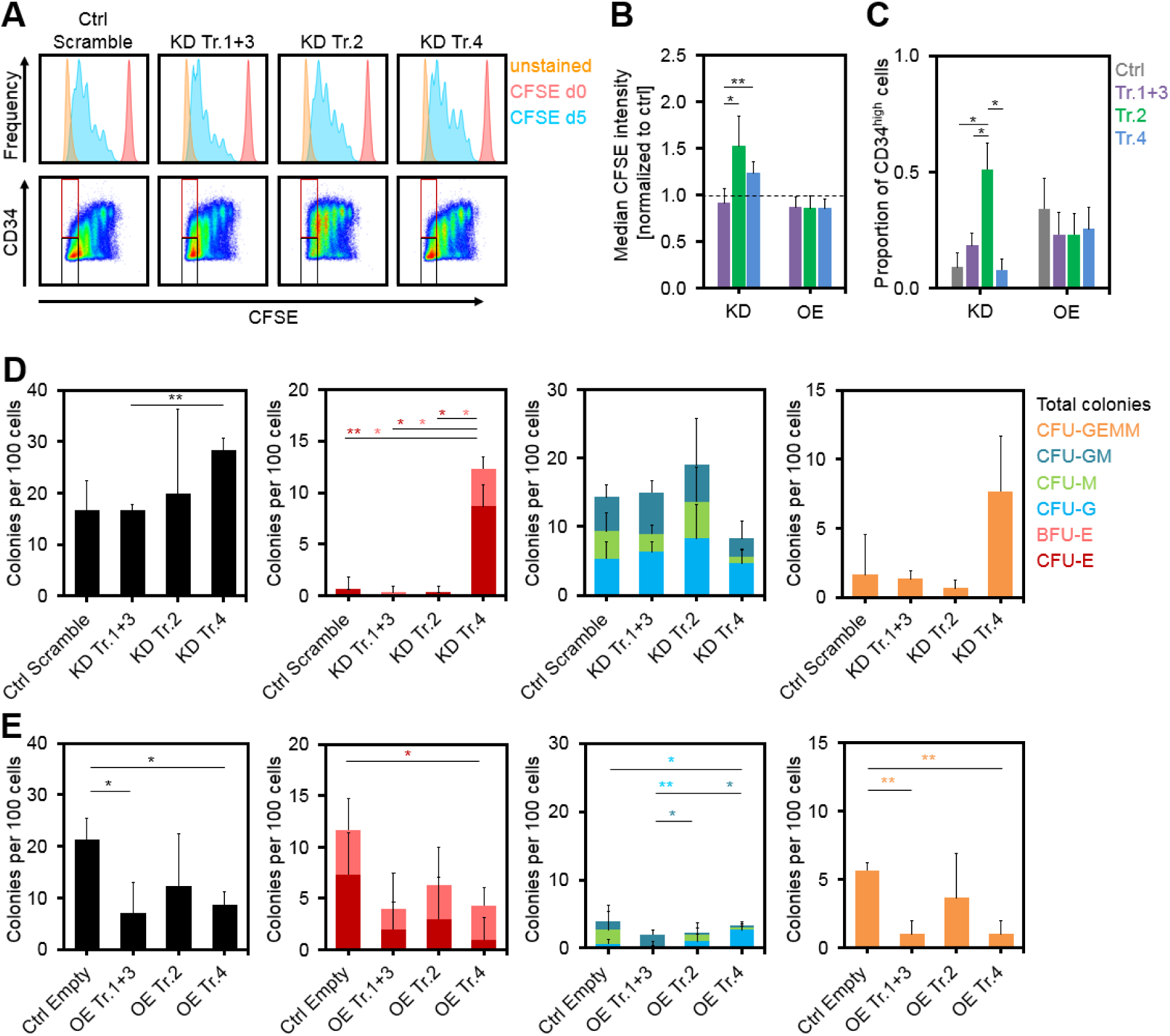
Impact of *DNMT3A* variants on proliferation and differentiation of HSPCs. (*A*) Histograms of residual CFSE staining to estimate proliferation of CD34 ^+^ cells after transfection with shRNAs after five days (blue). For comparison, measurements at day of HSPC isolation (day zero, no cell division, shown in red) and unstained controls at day five (shown in orange) are provided. ( *B*) Knockdown of transcripts 2 and 4 resulted in higher CFSE retention (slower proliferation) than the control (dashed line; mean ± SD; *n* = 3). (*C*) The proportion of CD34 high cells in the fast proliferating fraction was increased upon knockdown of transcript 2 (gates are indicated in Fig. 3A; mean ± SD; n = 3). (*D,E*) Colony forming unit (CFU) frequency (per 100 initially seeded HSPCs) was analyzed after knockdown (*D*) and overexpression (*E*) of DNMT3A transcripts (mean ± SD; n = 3). * P < 0.05, ** P < 0.01 (Student´s t-test).

Subsequently, we analyzed if modulation of *DNMT3A* variants impacts on maintenance of the surface marker CD34, as a surrogate marker for HSPCs. Downregulation of transcript 2 maintained CD34 expression even in the fraction of fast proliferating cells with more than five cell divisions (*P* < 0.05; *n* = 3; Figs. 3A,C). On the other hand, overexpression of the *DNMT3A* transcripts reduced the proportion of CD34^+^ cells in the fast proliferating fraction (Fig. 3C). In fact, differential CD34 expression was also observed in additional independent replicates, as well as corresponding DNAm and gene expression changes, indicating that DNMT3A2 is particularly relevant for loss of CD34 expression during culture expansion of HSPCs (Supplemental Fig. S2).

To investigate if *DNMT3A* variants affect colony forming unit (CFU) potential, we seeded HSPCs in methylcellulose at day five after infection with shRNAs or overexpression and analyzed colonies after two additional weeks of culture. Surprisingly, colonies of the erythroid lineage were significantly increased upon knockdown of transcript 4 (Fig. 3D) and reduced upon overexpression of transcript 4 (Fig. 3E). Thus, transcript 4 seems to impact on lineage-specific hematopoietic differentiation, although it does not comprise the functional methyltransferase domain. It is conceivable that effects of this isoform are mediated by other epigenetic or transcriptional modifiers by binding to the DNA with the N-terminal region (Suetake et al. 2011). Posttranslational modification of the *Dnmt3a* N-terminus was also shown to facilitate additional interactions (Chang et al. 2011). Overall, our data suggest that specific *DNMT3A* variants have different impact on proliferation and differentiation of HSPCs *in vitro*.

### Signatures of DNMT3A2 are recapitulated in acute myeloid leukemia

To determine if variant-specific signatures of DNMT3A are also coherently modified *in vivo* we utilized the AML dataset of The Cancer Genome Atlas (TCGA) (The Cancer Genome Atlas Research Network 2013). It has been demonstrated that AML cells with the R882H mutation have severely reduced *de novo* methyltransferase activity and focal hypomethylation at specific CpGs (Russler-Germain et al. 2014). Therefore, we reasoned that expression of *DNMT3A* transcripts in AML might also be reflected in our transcript-specific DNAm and gene expression signatures. As a surrogate for expression of individual transcripts we analyzed expression of transcript-specific exons, that were targeted by shRNAs in the knockdown experiments. In the subsequent analysis we have only observed significant correlations for the “*DNMT3A2*-exon” (ENSE00001486123), which might partly be attributed to the fact that the signatures for transcripts 1+3 and transcript 4 were smaller.

Initially, we analyzed if *DNMT3A* mutations in AML impact on the DNAm signature of DNMT3A2. To this end, we specifically focused on the CpGs that were differentially methylated upon knockdown of transcript 2 in HSPCs *in vitro*. AML samples with *DNMT3A* mutations revealed a significantly lower DNAm level in these DNMT3A2-associated CpGs than those without *DNMT3A* mutations (*P* = 0.03; Fig. 4A) and these samples were therefore excluded for further analysis. Notably, *DNMT3A2*-exon expression in AML patients revealed a highly significant correlation with DNAm levels at CpGs that were hyper- and hypomethylated upon knockdown of transcript 2 *in vitro* (*P* < 10^−100^ and *P* = 2*10^−37^, respectively; Fig. 4B). In analogy, *DNMT3A2*-exon expression in AML was also significantly correlated to expression of the 155 upregulated and the 70 downregulated genes of the *in vitro* transcript 2 signature (*P* = 6*10^−13^ and *P* = 0.0001, respectively; Figs. 4C,D). Furthermore, the average expression level of these 155 genes was significantly higher in those AML patients with below median expression of the *DNMT3A2*-exon (*P* < 0.0001; Fig. 4E). Similar results were observed when we used relative expression of the *DNMT3A2*-exon normalized by the overall *DNMT3A* expression level. Taken together, expression of *DNMT3A2*-exon is associated with variant-specific molecular signatures in AML.

Next, we analyzed if expression of transcript 2 might also be of clinical relevance in AML. In fact, expression of the *DNMT3A2*-exon was significantly lower in the AML sub-groups M4 and M5 of the French-American-British (FAB) classification (Fig. 4F). Furthermore, it was lower in patients with poor and intermediate cytogenetic risk score as compared to patients with favorable risk score (*P* < 0.01 and *P* < 0.001, respectively). Kaplan-Meier analysis (*P* = 0.019; Fig. 4G) and Cox regression analysis (*P* = 0.016) indicated that AML patients with lower expression of the *DNMT3A2*-exon have a significantly shorter overall survival. In comparison to the established molecular parameters for AML stratification, expression of *DNMT3A* variants has lower prognostic value, but our results support the notion that alternative splicing of *DNMT3A* is also relevant for the disease.

**Figure 4.**
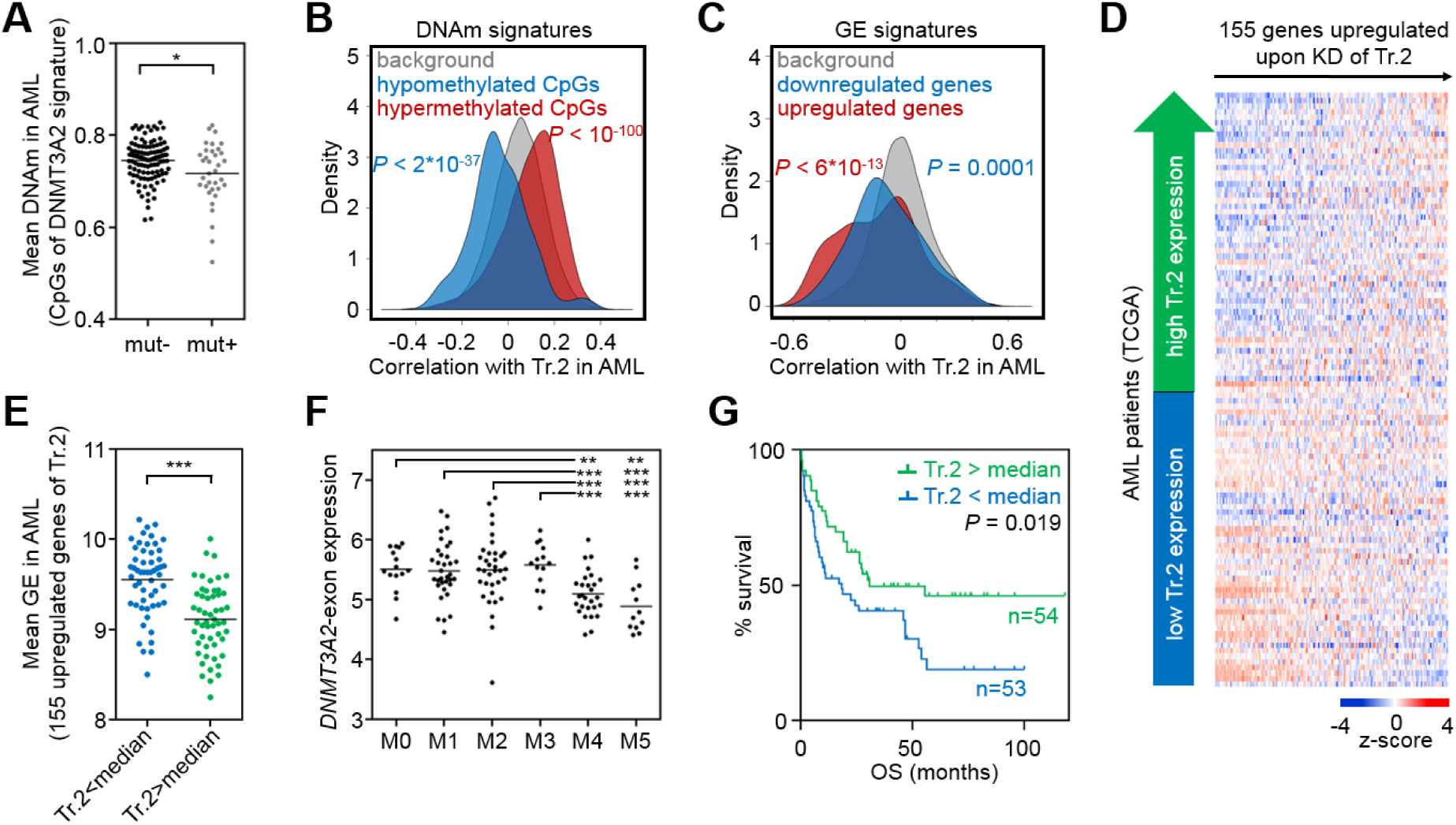
*DNMT3A2* signatures are coherently modified in acute myeloid leukemia. *DNMT3A2*-associated DNAm and gene expression signatures are recapitulated in the AML dataset of The Cancer Genome Atlas (TCGA) (The Cancer Genome Atlas Research Network 2013). (*A*) CpGs that revealed significant DNAm changes upon knockdown of transcript 2 in HSPCs *in vitro* had overall significantly lower mean DNAm levels in AML samples with a *DNMT3A* mutation– therefore samples with *DNMT3A* mutation were excluded for further analysis. (*B*) As a surrogate for transcript 2 expression we utilized the transcript-specific *DNMT3A2*-exon (ENSE00001486123). CpGs that were either hypo- or hypermethylated upon knockdown of transcript 2 in HSPCs *in vitro,* revealed overall highly significant correlation with expression of the *DNMT3A2*-exon in AML. (*C*) In analogy, genes that were differentially expressed upon knockdown of transcript 2 in HSPCs *in vitro* were significantly related to expression of the *DNMT3A2*-exon in AML patients. (*D*) Heatmap for association of *DNMT3A2*-exon expression in AML with expression of 155 genes that were upregulated upon knockdown of transcript 2 in HSPCs. (*E*) These 155 genes were on average significantly higher expressed in AML patients with lower *DNMT3A2*-exon expression (stratified by median). (*F*) *DNMT3A2*-exon is significantly lower expressed in AML subgroups M4 and M5 (FAB-classification). (*G*) Kaplan-Meier plot indicates that lower *DNMT3A2*-exon expression (stratified by median) is associated with shorter overall survival. * *P* < 0.05, ** *P* < 0.01, *** *P* < 0.001 (Mann W hitney test).

Targeting of DNMTs to specific sites in the genome is orchestrated by a complex interplay with other proteins, transcription factors, the histone code, and long non-coding RNAs (Yang et al. 2015; Kalwa et al. 2016). The results of this study add a new dimension to this complexity. The 27 exons of *DNMT3A* can be spliced into a multitude of different transcripts – although so far only five protein coding transcripts have been described. Our results demonstrate that different *DNMT3A* variants indeed have different transcript-specific molecular sequels that impact on hematopoietic differentiation and malignancy.

## Materials & Methods

### Cell culture of hematopoietic stem and progenitor cells

Umbilical cord blood (CB) was obtained after written consent according to guidelines approved by the Ethics Committee of RWTH Aachen Medical School (EK 187-08). CD34^+^ HSPCs were isolated from fresh CB using the CD34 Micro Bead Kit (Miltenyi Biotec, Bergisch-Gladbach, Germany) and cultured in StemSpan Serum-Free Expansion Medium (Stemcell Technologies, Vancouver, BC, Canada) supplemented with 10 μg/mL heparin (Ratiopharm, Ulm, Germany), 20 μg/mL thrombopoietin (PeproTech, Hamburg, Germany), 10 ng/mL stem cell factor (PeproTech), 10 ng/mL fibroblast growth factor 1 (PeproTech) and 100 U/mL penicillin/streptomycin (Lonza, Basel, Switzerland) (Walenda et al. 2011).

### Lentiviral knockdown and overexpression of *DNMT3A* variants

To knockdown *DNMT3A* transcripts we designed short hairpin RNAs (shRNAs) to target transcript-specific exons: exon 5 of transcripts 1+3 (ENSE00001486208), exon 2 of transcript 2 (ENSE00001486123), and exon 4 of transcript 4 (ENSE00001559474; Fig. 1A). In brief, forward and reverse oligonucleotides (Metabion, Planegg, Germany; Supplemental Table S3) were joined and ligated into the pLKO.1 vector (Addgene, Cambridge, MA, USA).

For constitutive overexpression, the *DNMT3A* transcripts were amplified from cDNA of human blood cells with the High-Capacity cDNA Reverse Transcription Kit (Applied Biosystems, Waltham, MA, USA) using various combinations of exon-specific primers (Supplemental Table S4) and cloned into the pLJM1-EGFP (Addgene) by replacing the EGFP gene. Successful cloning was validated by sequencing. About 200,000 HSPCs were infected one day after isolation and selected by treatment with puromycin (2.5 μg/mL; Sigma-Aldrich, St. Louis, MO, USA) at day two after infection.

### Quantitative RT-PCR

Knockdown or overexpression of *DNMT3A* variants was analyzed by real-time quantitative PCR (RT-qPCR) using the StepOneTM Instrument (Applied Biosystems). RNA was isolated at day 12 after infection, reverse transcribed, and amplified using the Power SYBR Green PCR Master Mix (Applied Biosystems) with transcript-specific primers (Supplemental Table S5). Gene expression was normalized to glyceraldehyde 3-phosphate dehydrogenase (GAPDH).

### DNA methylation profiles

DNA methylation profiles were analyzed for KD and OE condition, each in three independent biological replicates. Genomic DNA was isolated at day 12 after lentiviral infection with the NucleoSpin Tissue kit (Macherey-Nagel, Düren, Germany). For DNAm analysis we have chosen the Infinium HumanMethylation450 BeadChip (Illumina, San Diego, CA, USA) that covers about 480,000 representative CpG sites at single base resolution (including 99% of RefSeq genes and 96% of CpG islands) (Bibikova et al. 2011). In comparison to genome-wide analysis with reduced bisulfite sequencing data each of these CpGs is detected in all samples with relatively precise estimates of DNAm levels. Furthermore, this microarray platform enabled straight forward comparison with datasets of TCGA.

For further analysis of DNAm profiles, we excluded CpGs in single nucleotide polymorphisms and CpGs with missing values in several samples. Significance of DNAm was calculated in R using limma paired *t* test (adjusted for multiple testing; *P* < 0.05). Association of differentially methylated regions (DMRs) with the histone code was analyzed in ChIP-seq data from cultured CD34^+^ as provided by the IHEC portal (http://epigenomesportal.ca/ihec/). To estimate differences in levels of histone modifications (active: H3K4me3; enhancer: H3K4me1; repressive: H3K27me3 and H3K9me3), we calculated the average enrichment of the certain histone mark by a 250 bp window around DMRs (normalized by quantile normalization) as compared to the coverage of all CpGs on the microarray (Mann-Whitney test followed by multiple test correction). Enrichment of CpGs in relation to CpG islands (CGIs) or gene regions was based on the Illumina annotation and significance was estimated by hypergeometric distribution. Enrichment of short, core DNA-binding motifs of transcription factors within 100 bp around the differentially methylated CpG sites was performed in Python using the Regulatory Genomics Toolbox package (http://www.regulatory-genomics.org/motif-analysis/). Gene Ontology analysis was performed with the GoMiner software (http://discover.nci.nih.gov/gominer/) of genes associated with differentially methylated CpGs located in promoter or 5’UTR regions. Enrichment of specific categories was calculated by the one-sided Fisher’s exact *P*-value using all genes represented on the array as a reference.

### RNA expression profiles

Gene expression profiles were analyzed for KD and OE condition, each in three independent biological replicates. Total RNA was isolated at day 12 after infection with the NucleoSpin RNA kit (Macherey-Nagel) and analyzed with the Affymetrix Human Gene ST 1.0 platform (Affymetrix, Santa Clara, CA, USA). Raw data was normalized by RMA (Affymetrix Power Tools). Differentially expressed genes were filtered by at least 1.5-fold differential mean expression levels and adjusted *P* < 0.05, which were calculated in R using limma paired *t*-test. Gene Ontology analysis was performed with the GoMiner software (http://discover.nci.nih.gov/gominer/).

### Flow cytometric analysis

Carboxyfluorescein N-succinimidyl ester (CFSE) was used to monitor the number of cell divisions (Walenda et al. 2010). The immunophenotype of HSPCs was examined at day five after infection by staining with CD13-PE (clone WM15), CD34-APC (clone 581), CD38-PE (clone HIT2), CD45-V500 (clone HI30), CD56-PE (clone B159; all by Becton Dickinson, Franklin Lakes, NJ, USA), CD33-APC (clone AC104.3E3), CD133/2-PE (clone AC141; both Miltenyi Biotec) with or without CFSE staining. Cells were analyzed using a FACS Canto II (BD) with FACS Diva software (BD).

### Colony Forming Unit (CFU) assay

HSPCs were infected with lentivirus and selected with puromycin as described above and expanded for five days prior to CFU assay. Subsequently, 100 cells per well were seeded in methylcellulose based medium (HSC-CFU lite with EPO; Miltenyi Biotec). Granulocyte (CFU-G), macrophage (CFU-M), granulocyte/macrophage (CFU-GM), erythroid (BFU-E and CFU-E), and mixed colonies (CFU-GEMM) were counted according to manufacturer’s instructions after 14 days (Walenda et al. 2011).

### Analysis of AML datasets

DNA methylation and gene expression signatures of *DNMT3A* variants were further analyzed in acute myeloid leukemia (AML) patients of The Cancer Genome Atlas (TCGA) portal (The Cancer Genome Atlas Research Network 2013). We focused on 142 patients with data available for HumanMethylation450 Bead Chips, RNA sequencing, and whole exome sequencing. For comparison of DNAm data of the DNMT3A2 signature only 7,074 CpGs (6,747 hyper- and 327 hypomethylated) of the 8,905 significant differentially methylated CpGs were provided by the TCGA repository. Raw gene expression data was normalized using the variance stabilizing transformation. Expression levels of transcript-specific exons (ENSE00001486208 for transcripts 1+3; ENSE00001486123 for transcript 2, ENSE00001559474 for transcript 4) were used as a surrogate for corresponding transcript expression. Alternatively, we normalized expression of transcript-specific exons to the mRNA level of *DNMT3A* and these relative expression levels provided similar results. Correlations with the signatures were calculated in R. For survival analysis, we used Kaplan-Meier (K-M) and Cox proportional hazards model. For K-M analysis, patients were stratified into two groups according to the median expression level of the *DNMT3A2*-exon. We defined overall survival (OS) as the survival time from the first day of diagnosis to day of death by any cause. K-M plots and Mann-Whitney statistics were generated with GraphPad Prism 6.05 (GraphPad Software, San Diego, CA, USA) and Cox proportional hazard model was calculated in R.

## Acknowledgments

This work was supported by the Interdisciplinary Center for Clinical Research (IZKF) within the Faculty of Medicine at the RWTH Aachen University (O1-1), by the Else Kröner-Fresenius-Stiftung (2014_A193), by the German Research Foundation (WA 1706/8-1), and by the German Ministry of Education and Research (01KU1402B).

## Author Contributions

T.B., J.F. and A.R. designed the study, performed experiments, and analyzed the data. S.H.H. generated DNAm and gene expression profiles. F.T., C.-C.K. and I.G.C. conducted bioinformatic analysis. T.G. provided cord blood samples. E.J., M.Z. and W.W. conceived the study, participated in data analysis, and coordinated research. T.B., J.F. and W.W. wrote the manuscript and all authors read, edited, and approved the final manuscript.

## Disclosure Declaration

RWTH Aachen Medical School has applied for a patent on the *DNMT3A* epimutation. Apart from this the authors do not have a conflict of interest to declare.

